# Nucleation-dependent aggregation kinetics of Yeast *Sup*35 fragment GNNQQNY

**DOI:** 10.1101/2020.07.27.221150

**Authors:** Gunasekhar Burra, Mahmoud B. Maina, Louise C. Serpell, Ashwani K. Thakur

## Abstract

An N-terminal hepta-peptide sequence of yeast prion protein Sup35 with the sequence GNNQQNY serves as an ideal model for structural understanding of amyloid assembly and kinetics. In this study, we used a reproducible solubilisation protocol that allows the generation of homogenous monomeric solution of GNNQQNY to understand the molecular details of its self-assembly mechanism. The aggregation kinetics data show that the GNNQQNY sequences follow nucleation-dependent aggregation kinetics with a critical nucleus of size ~7 monomers and that the size and efficiency of nucleation was found to be inversely related to the reaction temperature. The generated nucleus reduces the thermodynamic energy barrier by acting as a template for further self-assembly and results in highly ordered amyloid fibrils. The fibers grown at different temperatures showed similar Thioflavin T positivity, Congo red binding and β-sheet rich structures displaying a characteristic cross-β diffraction pattern. These aggregates also share morphological and structural identity with those reported earlier. The mature GNNQQNY fibers exerted no significant oxidative stress or cytotoxicity upon incubating with differentiated SHSY5Y cells. To our knowledge, this is the first study to experimentally validate previous predictions based on theoretical and molecular dynamics simulations. These findings will provide the basis for understanding the kinetics and thermodynamics of amyloid nucleation and elongation of amyloidogenic systems associated with many systemic and neurodegenerative diseases.

## Introduction

Synthetic peptide fragments that mimic the formation of amyloid fibrils from full-length amyloidogenic proteins are widely used to understand the physico-chemical aspects of protein aggregation under *in vitro* conditions.^*1, 2*^ The smaller length peptides possess relatively lower complexity thereby providing an ideal system for *in vitro* studies.^*3–5*^ A seven residue sequence ‘GNNQQNY’ spanning from residue 7 to 13 of full length Sup35, a eukaryotic prion protein, serves as an ideal model for understanding the structural aspects of amyloids since the crystal structure has been solved in two forms.^*6*^ Furthermore, GNNQQNY can trigger the formation of fibril-core in yeast Sup35 prion fibrils.^*3*^ In yeast, Sup35 protein plays a major role in translation termination and can undergo prion-like structural conversion. Unlike other infectious and toxic prion proteins, prions formed by Sup35 are non-toxic and are stably inherited for many cellular generations, thus resulting in a characteristic phenotypic trait.^*7–10*^ This is due to functionally inactive state of Sup35 in prions leading to impaired translation termination.^*7–10*^

X-ray fibre diffraction from GNNQQNY amyloid fibrils shows a characteristic meridional reflection at ~4.7 Å and an equatorial reflection at ~10 Å, indicating the inter-strand (backbone) and inter-sheet (side chain) distances.^*11–13*^ These diffraction signals arise from the characteristic cross-β spine containing steric zipper structure in which the β-strands run perpendicular to the long fiber axis.^*14, 153, 14, 16*^ GNNQQNY fibrils are long, unbranched and display *apple-green* birefringence upon binding to Congo red and *green-yellow* fluorescence in solution with thioflavin T.^*14*^

Previous studies based on molecular dynamic simulations and theoretical predictions have shown that GNNQQNY undergoes nucleation-dependent aggregation kinetics through which it forms fibrils rich in parallel β-sheets.^*2, 14, 15, 17, 18*^ This involves the formation of a metastable critical nucleus within the monomer pool along the path of aggregation. Once the nucleus is formed, further monomer addition occurs at a much faster rate leading to the elongation of amyloid fibrils.^*19*^ Using structure-based theoretical energetics, Nelson *et al* (2005) predicted that three-to-four GNNQQNY molecules constitute the metastable nucleus formed during the aggregation.^*14*^ A different study based on molecular dynamic simulations suggested a nucleus of size about four-five monomers and five-six monomers at 280 K and 300 K, respectively.^*18*^ A more recent study by Langkilde *et al* (2015) based on small angle X-ray scattering (SAXS) and thioflavin T analyses suggested the lack of intermediates during the fibrillation process and suggested that the elongation process occurs most likely *via* monomer addition.^*20*^ Although significant advancement has been achieved in understanding the molecular structure of amyloid fibers formed by GNNQQNY, the *in vitro* kinetics and thermodynamics associated with its formation is yet to be elucidated.

Here, we used a solubilisation protocol that allows the generation of homogenous monomeric solution reproducibly. This enabled the study of aggregation and nucleation kinetics analyses by quantitative RP-HPLC-based sedimentation assay. The sedimentation, thioflavin T binding, right-angle light scattering and seeding kinetics analyses results have experimentally revealed that the GNNQQNY sequence follows nucleation-dependent aggregation kinetics. The nucleation kinetics analysis indicates the formation of a critical nucleus of size seven monomers that acts as a template to drive the aggregation reaction under physiological conditions (pH 7.4 and 37 °C). The effect of temperature on aggregation and nucleation was also evaluated. This study provides experimental proof for the previous predictions which are based on theoretical and molecular dynamic simulations. The mature GNNQQNY fibrils generated did not result in a significant increase in cytotoxicity or cell death.

## Materials and Methods

### Materials

Uncapped GNNQQNY (95% pure), Trifluoroacetic acid (TFA), formic acid, Congo red and thioflavin T were purchased from Sigma Aldrich. Uncapped PGQ_9_I (K_2_-Q_9_-PG-Q_4_-I-Q_4_-PG-Q_9_-PG-Q_9_-K_2_) and PGQ_9_A (K_2_-Q_9_-PG-Q_4_-A-Q_4_-PG-Q_9_-PG-Q_9_-K_2_) peptides were obtained from Keck Biotechnology Centre, Yale University in crude form and were then purified to near homogeneity. These model peptides are used to understand the aggregation mechanism of polyglutamine-containing sequences.^*21, 22*^ HPLC-grade acetonitrile and HCl were procured from Merck and sodium-azide was from SD fine chemicals Ltd. 500 mL of sterile 10X Phosphate-buffered saline (PBS) was prepared by dissolving NaCl (40 g), KCl (1 g), Na_2_HPO_4_ (7.2 g) and KH_2_PO_4_ (1.2 g) procured from Merck as per Cold Spring Harbor protocols.

### Solubilization of peptides

GNNQQNY peptide (2 mg/mL) was solubilised in milliQ-water acidified to pH 2.0 using HCl. The resulting solution was gently swirled to ensure complete solubilisation of the peptide and ultracentrifuged for 2 h at 80, 000 rpm and 25 °C. The top 2/3^rd^ of the supernatant was collected carefully without disturbing the bottom 1/3^rd^ of the solution that may contain the pre-aggregated peptide and used for further studies. PGQ_9_I and PGQ_9_A peptides were solubilised using established protocols.^*22–26*^

### Mass spectrometry

5 μL of GNNQQNY peptide solubilized in water-HCl (pH 2.0) was injected into time-of-flight (TOF) mass analyzer provided with electrospray ionization (ESI) source (Waters Q-TOF premier HAB213). The *m/z* spectrum recorded at a mass range of 50–3500 Da was plotted using OriginPro 8.5 data analysis and graphing software and the purity was estimated by analyzing the observed molecular ionization states of the peptide.^*24*^

### Analytical size–exclusion chromatography (SEC)

The GNNQQNY peptide solubilized in water-HCl (306 μM) was adjusted to pH 7.4 by using 10X PBS such that the final concentration of PBS in the reaction mixture becomes 1X. 100 μL of this peptide solution was subjected to SEC analysis by passing through Superdex peptide 10/300 GL column (GE healthcare) connected to Bio-Rad (Biologic Duoflow) made fast protein liquid chromatography (FPLC) system. The elution profile of the peptide was monitored in 1X PBS (pH 7.4) at a flow rate of 0.5 mL/min. The absorbance of the peptide was recorded at 215 nm under a constant pressure of about 80 psi.^*24, 25*^

### Sedimentation assay

The aggregation was monitored in PBS (pH 7.4) at 37 °C, 23 °C and 4 °C. Sodium azide (0.05%) was added to prevent microbial contamination and growth.^*22, 27–29*^ The aggregation kinetics was monitored by measuring the decline in monomer concentration as a function of time. At regular time intervals, an aliquot of sample from the ongoing aggregation reaction was subjected to ultracentrifugation for 30 min at 25,000 rcf and 25 °C. The supernatant was collected, adjusted to 20% formic acid and then injected into RP-HPLC to determine the monomer concentration based on the predetermined standard curve (See SI).^*30, 31*^ Adding formic acid has been shown to prevent aggregation in the vial by reducing the pH of the reaction.^*27*^

### Light scattering (LS) and Thioflavin T (ThT) assays

These assays were performed as described previously by taking 120 μL of ongoing aggregation reaction sample in a Quartz SUPRASIL Ultra-micro cell.^*25*^ The intensity of light scattered by the sample was determined by setting the emission and excitation wavelengths at 450 nm and their slit widths at 2.5 nm. ThT fluorescence intensity was measured by adding ThT stock to a final concentration of 100 μM to the above sample. This was scanned by resetting the excitation wavelength to 450 nm (slit width, 5 nm) and the emission wavelength to 489 nm (slit width, 5 nm). These assays were carried out on a LS 55 spectrofluorimeter (Perkin Elmer). The blank was subtracted from consecutive spectra which were averaged to obtain the final LS and ThT spectra.

### Seeding assay

Seeding assay was performed as described previously.^*21, 22*^ Seeding reactions were monitored in PBS (pH 7.4) at 37 °C by adding the preformed seeds (2%, wt/wt) grown at similar conditions. No preformed seeds were added for the control reactions used for comparison. The kinetics of aggregation was determined by measuring the concentration of monomers at different time intervals using sedimentation assay. The data was plotted between monomer percentages and time (h) for comparison.

### Nucleation Kinetics analyses

Different concentrations of freshly disaggregated GNNQQNY were prepared in PBS at pH 7.4 and incubated at 37 °C and 23 °C. The concentration of the soluble monomer in the ongoing reactions was measured at different time points using sedimentation assay. Monomer concentration (μM) versus time^2^ (s^2^) graphs for each concentration were plotted by taking as many data points as possible within the initial 20% drop in monomer concentration. The slopes from these time^2^ graphs were used to construct log[initial concentration (M)] versus log[slope] plots. The slope obtained upon subjecting this plot to linear curve fit was used to solve the equation, slope = n* + 2, where n* is the size of the critical nucleus.^*19, 21, 22, 27, 32*^

### Thioflavin T (ThT) and Congo-red staining of fibrils

Working standard solutions of 0.8 mg/mL of ThT and 1 mg/mL of Congo red solubilised in MilliQ-water and filtered using 0.2 μ MDI syringe filter were used to stain the mature amyloid fibers of GNNQQNY. 10 μL of peptide solution was spread on a fresh glass slide and allowed to air-dry. The dry peptide film was then stained with a drop of working standard solution of ThT and Congo-red; incubated for 30 min, excess dye was removed by washing with MilliQ-water and subsequently air dried for 30 min. Bright field images and the corresponding ThT fluorescence and Congo-red birefringence images were captured using appropriate light filters on Leica DM2500 fluorescent microscope.^*3, 33*^

### Fourier-transform infrared (FTIR) spectroscopy

FTIR spectroscopy analysis was carried out using Bio-ATR II cell mounted on Tensor 27, Bruker FTIR instrument.^*22, 25, 34*^ GNNQQNY aggregates were subjected to washing thrice by using milliQ-water and then 50 μL was loaded on to the Bio-ATR II cell. A total of 120 scans at a resolution of 4 cm^−1^ under constant purging of nitrogen gas were collected at room temperature and averaged for the final FTIR spectra. The obtained spectra were auto-corrected for water vapour and the buffer. The resultant primary spectrum was then subjected to second derivative analysis using nine-point smoothing by Opus 7.2 spectroscopy software (Bruker Optics).

### Transmission electron microscopy (TEM)

Samples for electron microscopy analysis were prepared by inverting the carbon-coated side of TEM grid on 5 μL of aggregate sample, incubated for thirty seconds and the excess sample was carefully blotted by using filter paper.^*22, 25, 35*^ It was rinsed with 10 μL of milliQ-water and then negatively stained by inverting on 5 μL of 2% uranyl acetate for 5-10 seconds. The stained grids were left overnight to air-dry and then imaged using a FEI-Tecnai G2 12 Twin (120 KV) transmission electron microscope.

### X-ray fiber diffraction

200 μL peptide stock solution (10 mg/mL) was incubated at 37 °C, 23 °C and 4 °C for several days to generate mature fibers. The fibers were then aligned by placing 20 μL of each sample between two wax-filled capillaries and allowed to air-dry overnight. The aligned fibers were placed on a goniometer head mounted on a Rigaku X-ray diffraction machine (Sevenoaks, UK) containing a rotating anode (CuKα) and Raxis IV++ detector. The data was recorded using specimen-to-film distance of 50 mm or 100 mm.^*36*^

### Cytotoxicity and Cell viability assays

Undifferentiated SHSY5Y Neuroblastoma cells were differentiated using established protocol.^*37, 38*^ The undifferentiated cells (ATCC CRL-2266™) were cultured at 37 °C and 5% CO_2_ incubator maintained in Dulbecco’s Modified Eagle Medium: Nutrient Mixture F-12 (DMEM/F-12) (Life Technologies, United Kingdom), mixed with 1% (v/v) L-glutamine (L-Gln) (Invitrogen), 1% (v/v) penicillin/streptomycin (Pen/Strep) (Invitrogen) and 10% (v/v) Fetal Calf Serum (FCS) (henceforth called media). The undifferentiated cells were seeded to 60% confluency to adhere for 24 h in a CellCarrier-96 Ultra Microplates (Perkin Elmer), which was followed by the media replacement with a fresh media containing FCS reduced to 1% and supplemented with 10 μM trans-Retinoic acid (Abcam) for 5 days. A media exchange with fresh media supplemented with the retinoic acid was done after 2 days. At day 5, the cells were incubated with a FCS-free media supplemented with 2 nM brain-derived neurotrophic factor (BDNF) (Merck Millipore). After 2 days, the media was replaced with FCS and BDNF free media and the cells were treated with fully grown amyloid fibers from GNNQQNY (5 μM and 20 μM) and PGQ_9_A (5μM and 20 μM) peptides for 48 hours. Phosphate buffered saline containing no aggregates were used as negative control, whereas 1 mM H_2_O_2_ was used as positive controls. Towards the end of the incubation period, the cells were treated for 45 min with 5 μM CellROX Green Reagent to evaluate the level of oxidative stress (C10444, Life technologies UK). This was followed by incubation for 15 min with ReadyProbes reagent (Life Technologies) to evaluate cell death. The cells were imaged using Operetta CLS high-content imaging system (PerkinElmer) using DAPI, FITC and TRITC filters at 37 °C and 5% CO_2_. A minimum of three independent experiments each containing at least 5000 cells were analysed using the Harmony software automated analysis algorithm within the system to ensure the reproducibility of the findings.

## Results and discussion

### Solubilisation and characterization of GNNQQNY

Previous studies on GNNQQNY aggregation have been unable to unambiguously resolve the kinetics of aggregation due to contamination with pre-aggregated peptide and most studies have relied on in-silico analysis.^*17, 18*^ Here, we used an acid solubilisation and centrifugation procedure that removed pre-aggregated peptide and resulted in monomeric solution. SEC of GNNQQNY showed a single peak suggesting the homogenous nature of the solution (Fig. 1A). The elution of GNNQQNY (836.8 Da) after PGQ_9_I (1974.4 Da) but prior to sodium azide (NaN_3_, 65 Da) further confirmed the absence of higher order structures in the solubilised fraction (Fig. 1A). This fraction when subjected to ESI-MS showed a characteristic *m/z* value of 836.4 Da, thus confirming the identity of GNNQQNY monomeric solution (Fig. 1B). Presence of only a single peak corresponding to the monomer (836.4 Da) also confirms the purity (>95%) of the peptide (Fig. 1B).

**Fig. 1:**
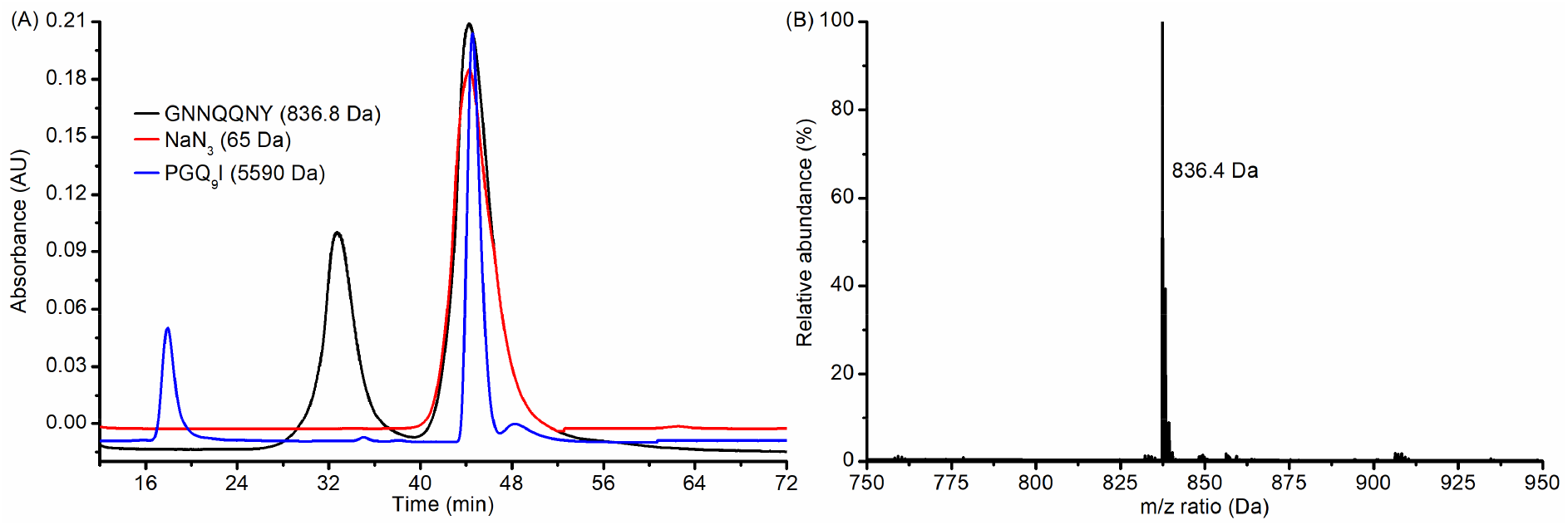
Characterization of soluble monomers of GNNQQNY. Size-exclusion chromatography analysis of GNNQQNY monomer in comparison with sodium azide (NaN_3_) and PGQ_9_I peptide (A). ESI-MS analysis confirming the monomeric (836.4 Da) state of the soluble fraction (B).

These data established that water-HCl (pH 2.0) solubilises GNNQQNY peptide and allows for initiating the aggregation reaction. This is probably due to the low pH (2.0) which is well below the isoelectric point (pI) of GNNQQNY (5.52). At low pH, the GNNQQNY molecules are expected to attain an overall positive surface charge. The presence of same charge at the peptide surface promotes its interaction with water, rather than with other peptide molecules thereby making it more soluble.^*39*^ Henceforth this procedure has been followed to obtain the monomeric solution needed for carrying out further experiments in this study.

### Aggregation kinetics

The aggregation kinetics was determined by measuring the monomer concentration at different time intervals based on the standard curve generated using Kuipers and Gruppen method (Fig. S1).^*40*^ Kinetics based on sedimentation assay at 1253±12 μM showed a clear lag-phase (nucleation-phase) for about 12 h of incubation during which the drop in the monomer concentration was found to be negligible (Fig. 2A, Black curve). Also, it took about 25 h for the initial 20% drop in monomer, during which the actual critical nucleus is believed to form in classical nucleation-dependent aggregation mechanism.^*27, 32, 41, 42*^ Together, this data suggests a lag-phase of about 20-25 h for GNNQQNY incubated at an initial concentration of 1253±12 μM, during which an energetically unfavourable species called nuclei are formed.

**Fig. 2:**
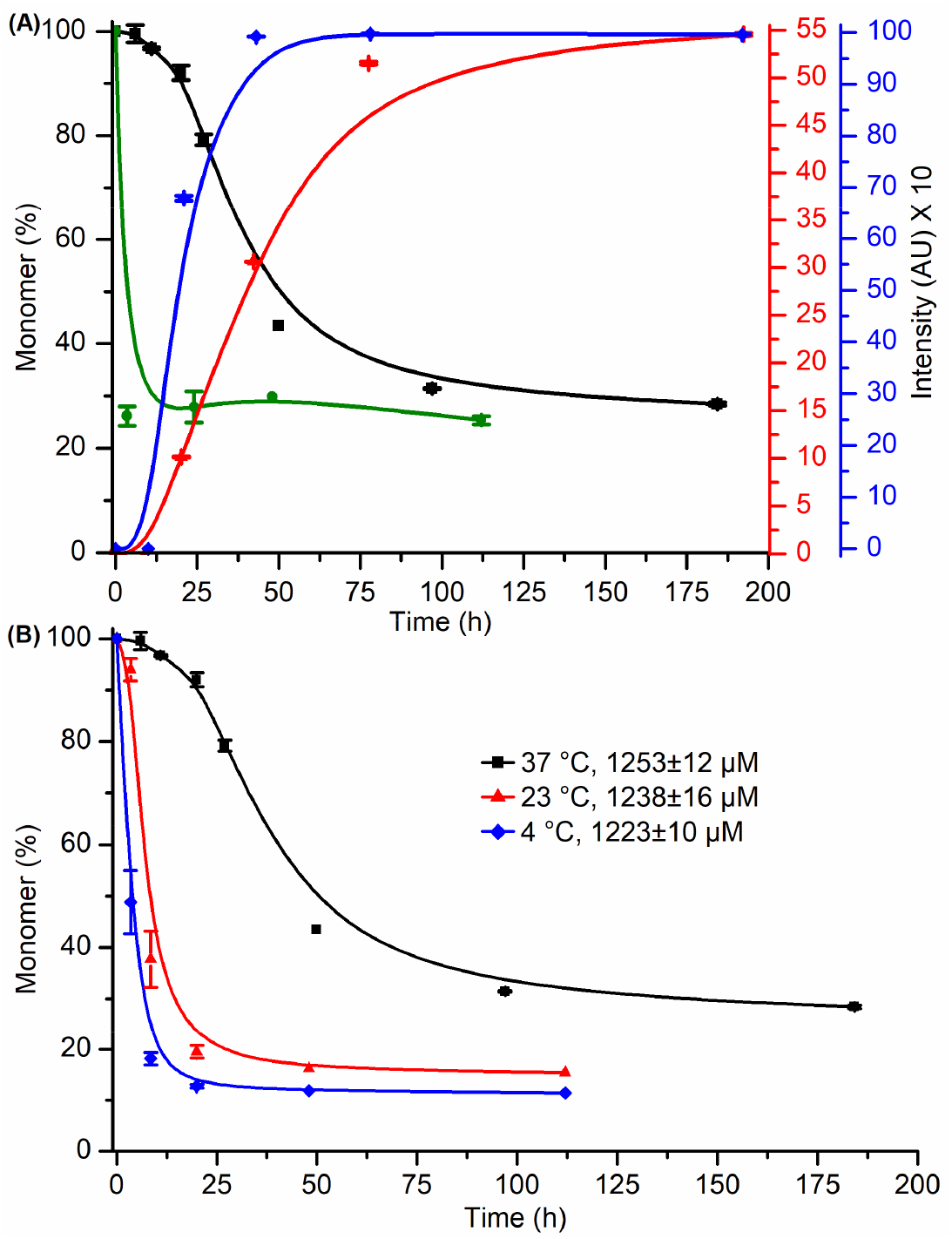
Aggregation kinetics analysis of GNNQQNY peptide. (A) Aggregation kinetics analyses at pH 7.4 and 37 °C based on sedimentation (Black), light scattering (Blue), ThT (Red) and seeding (Olive green) assays showing nucleation-dependent aggregation. (B) Shows the temperature-dependent aggregation kinetics. Error bars indicate standard deviation calculated from triplicate reactions.

The formation of this high energy species (nucleus) is followed by a rapid decrease in monomer concentration, indicating an elongation-phase. This finally reached an equilibrium or stationary-phase where no further drop in monomer concentration was observed (Fig. 2A, Black curve). The concentration of monomer at the stationary phase is also termed as critical concentration (C_r_), at or below which the efficiency of nucleation is almost negligible.^*19, 27, 32, 43*^ The initial delay in aggregation (lag-phase/nucleation-phase) was due to highly unfavourable equilibria that raise the thermodynamic energy barrier for polymerisation. The peak of the energy barrier that corresponds to the transient critical nucleus species makes the following downstream steps thermodynamically favourable, thus promoting the elongation.

The other characteristic features of a classical nucleation-dependent aggregation kinetics includes; (1) the elimination of lag-phase in presence of preformed seeds or aggregates, (2) lack of ThT binding until the nucleation of β-strands/β-sheets, and (3) the dependence of rate of aggregation on the initial reaction concentration.^*27, 43*^ These characteristics were tested by using appropriate methods to confirm the observations made using sedimentation assay. As expected, the addition of preformed seeds (2%, wt/wt) greatly enhanced the rate of aggregation of the same concentration of monomer by spontaneously reducing the monomer concentration to about 25% in just 3-4 h of incubation (Fig. 2A, Olive green curve). This confirms the abrogation of nucleation-phase (lag-phase), which is primarily because the added aggregates reduce the thermodynamic energy barrier by acting as templates for monomer addition and thus drives the aggregation via the elongation phase.^*22, 27, 41–43*^ Indeed, this supports the templated aggregation hypothesis reported in earlier study where the introduction of *in vitro* generated amyloid seeds of recombinant Sup35 protein can induce aggregation in cells.^*44*^

Specific binding of ThT dye to assembled structures results in enhanced fluorescence at 489 nm.^*45, 46*^ However, enhancement in fluorescence of ThT was not observed up to 12 h of incubation, confirming the absence of ThT positive structures throughout the nucleation-phase (Fig. 2A, Red curve). Fluorescence enhancements during the elongation-phase lead to the saturation after reaching the equilibrium-phase. Thus, the kinetics monitored using ThT has complemented the kinetics observed using sedimentation assay (Fig. 2A). The observation of a similar trend even by light scattering assay which indicates the formation of higher-order structures over time further confirmed the data obtained by the above techniques (Fig. 2A, Blue curve). An increase in ThT fluorescence associated with a decrease in monomer concentration after the lag-phase provides much stronger evidence towards the nucleation-dependent aggregation mechanism (Fig. 2A).

The concentration–dependence of GNNQQNY aggregation was established by monitoring the aggregation kinetics at six different staring concentrations. An increase in the lag-phase was observed upon reducing the starting concentration of the reaction (Fig. 3A). This inverse relationship confirms that the efficiency of nucleation decreases upon reducing the starting concentration thereby increasing the lag-phase.^*2, 19, 27, 32, 42, 43*^ All these *in vitro* observations confirmed the nucleation-dependent aggregation kinetics for GNNQQNY and provides experimental piece of evidence for the speculations made using theoretical and molecular dynamic simulation studies.^*6, 14, 15, 17, 18*^

**Fig. 3:**
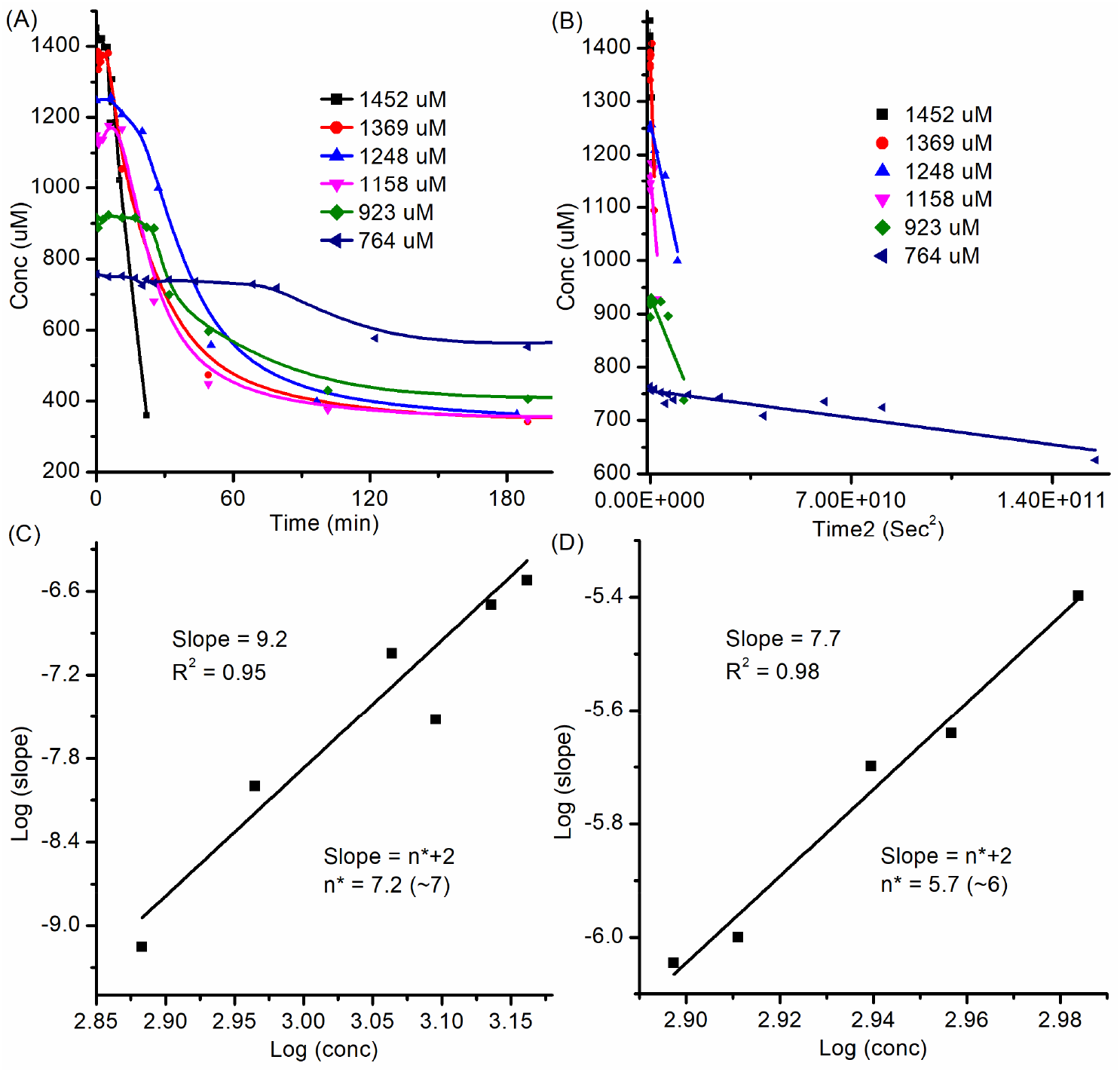
Nucleation kinetics analysis of GNNQQNY. (A-C) Shows the nucleation kinetics at physiological conditions of pH 7.4 and 37 °C. Concentration-dependent aggregation kinetics (A) from which the initial 20% drop in the monomer concentration was plotted against time^2^ (s^2^) to generate linear time^2^ plot (B). The log of slope obtained from (B) for each line plot was plotted against the log of respective initial concentration to obtain the log-log plot (C). (D) Represents the log-log plot for the nucleation kinetics at pH 7.4 and 23 °C.

However, this study deviates from the recent study by Langkilde *et al* (2015) where no structural nucleus was observed during the fibrillation process and it was suggested that the elongation process occurs most likely *via* monomer addition.^*20*^ In their study, the initial monomer concentration as high as 10.3 mM and 10.4 mM were used for the analysis.^*20*^ As any nucleation-dependent aggregation kinetics is governed by the initial monomer concentration, the free energy of transition of monomers to fibrils is strongly negative at very high initial concentration.^*14, 19, 27, 32, 42*^ This may be the explanation that no oligomer species or nucleus was observed. It is also possible that the nucleus formed at such high concentration is too small to be observed by SAX or too short lived. These speculations were further confirmed by measuring the size of the critical nucleus formed during aggregation.

### Size of critical nucleus

In order to measure the size of the transient critical nucleus (n*) formed on the path of GNNQQNY aggregation, aggregation reactions were set up at different initial concentrations (Fig. 3A). As many time points as possible were recorded for the first 20% drop in the monomer concentration and were used to construct the concentration (μM) versus time^2^ (Sec^2^) plots (Fig. 3B). The time (Fig. 3A) and time^2^ (Fig. 3B) plots have shown a clear concentration-dependence for the aggregation of GNNQQNY. The log of slope of each linear time^2^ plot was plotted against the respective log of initial concentration (M) to obtain a linear log-log rate-concentration plot with a slope of 9.2 (Fig. 3C). Substituting this value in the equation, slope= n*+2, resulted in n*=7.2 (Fig. 3C). This suggests a critical nucleus of size seven (n*=7, after round off), which means that seven monomers associate to form a highly unstable nucleus that drives the aggregation reaction forward. The size of the nucleus observed in this study is relatively larger (n*=7) when compared to the previously reported theoretical (n*=3-4) and simulation (n*=4-5 at 280 K and n*=5-6 at 300 K) based studies.^*14, 18*^ This is probably because the aggregation reaction in this study was monitored at physiological temperature (310 K) unlike the previous studies where a relatively lower temperature was used.

### Effect of temperature on aggregation

The effect of temperature on the nucleation event was investigated by carrying out a thorough temperature-dependent aggregation and nucleation kinetics analysis. Comparison of kinetics of aggregation monitored at 37 °C with that of 23 °C and 4 °C showed a reduction in the length of lag-phase upon reducing the temperature (Fig. 2B). Briefly, the time taken for initial 20% drop in monomer concentration was greatly reduced from ~25 h observed at 37 °C (Fig. 2B, Black curve) to ~5 h at 23 °C (Fig. 2B, Red curve) and ~75 min at 4 °C (Fig. 2B, Blue curve). In agreement with this observation, the nucleation kinetics analysis monitored at 23 °C resulted in a critical nucleus of size six (n*=6) monomers, which is relatively smaller than the nucleus (n*=7) observed at 37 °C (Fig. 3C, D). We have also monitored the nucleation kinetics at 4 °C but it was not possible to quantify the size of the critical nucleus due to rapid aggregation rate observation at this low temperature (data not shown). However, this suggests even a smaller nucleus at 4 °C as compared to 23 °C. Overall, these data suggest that the efficiency of nucleation increases thereby decreasing the lag-phase with decreasing the temperature of the aggregation reaction and supports the observations reported by Nasica-Labouze and Mousseau, 2012.^*18*^

### Molecular and structural characterization of the mature fibrils

The mature fibrils obtained at the end of the aggregation reactions monitored at different temperatures were stained using ThT and Congo red (CR). Using the bright field imaging, appropriate areas containing fibrils were chosen and exposed under fluorescein gate for ThT and polarized light for CR binding analysis (Fig. S2). Binding of ThT resulted in distinct bright *yellow-green* fluorescence all along the length of the fibrils (Fig. S2 A-C). Similarly, when the CR stained samples were subjected to polarised light microscopy, *apple-green* birefringence was observed along the fibrils (Fig. S2 D-F).^*47*^ The substantial enhancement of ThT fluorescence emission and CR birefringence property makes them particularly powerful and convenient tools for amyloid detection.^*33, 48*^ Previous studies have also shown that the binding of these dyes is associated with the presence of amyloid fibrils.^*33, 45, 46, 48–50*^ Thus, the use of ThT in conjunction with CR strongly suggests that the observed aggregates are amyloid fibrils by nature.

The amyloid structures of GNNQQNY were further confirmed by FTIR and TEM analyses. Infrared spectroscopy analysis showed a characteristic β-sheet band at 1631 cm^-1^ along with a corresponding minor band at 1690 cm^−1^, indicating their amyloidogenic nature (Fig. 4A). The other two bands represent the side chain NH_2_ defect (1605 cm^−1^) and C=O stretching (1670 cm^−1^) vibrations of Gln/Asn residues (Fig. 4A).^*25*^ Even, the aggregates generated through seeding reaction resulted in an IR spectrum that is similar to the aggregates generated at three different temperature conditions (Fig. 4A). TEM images of fibers showed long unbranched fibrous-to-crystalline structures similar to those reported earlier for GNNQQNY (Fig. 4B-D).^*20, 36, 47, 51*^ Furthermore, the X-ray fiber diffraction data showed the characteristic meridional (4.75 Å) and equatorial (9.25 Å) diffraction signals, thus confirming the cross-β architecture of the amyloid fibrils generated in this study (Fig. 5E-G).^*3, 14, 15, 36, 47, 51*^ From these observations, it is evident that the final aggregates generated at three different temperatures (37 °C, 23 °C and 4 °C) are similar to each other with respect to their molecular architecture and structural morphology. The only striking difference observed is the size of the nucleus that is proportional to the length of the lag-phase and the reaction temperature.

**Fig. 4:**
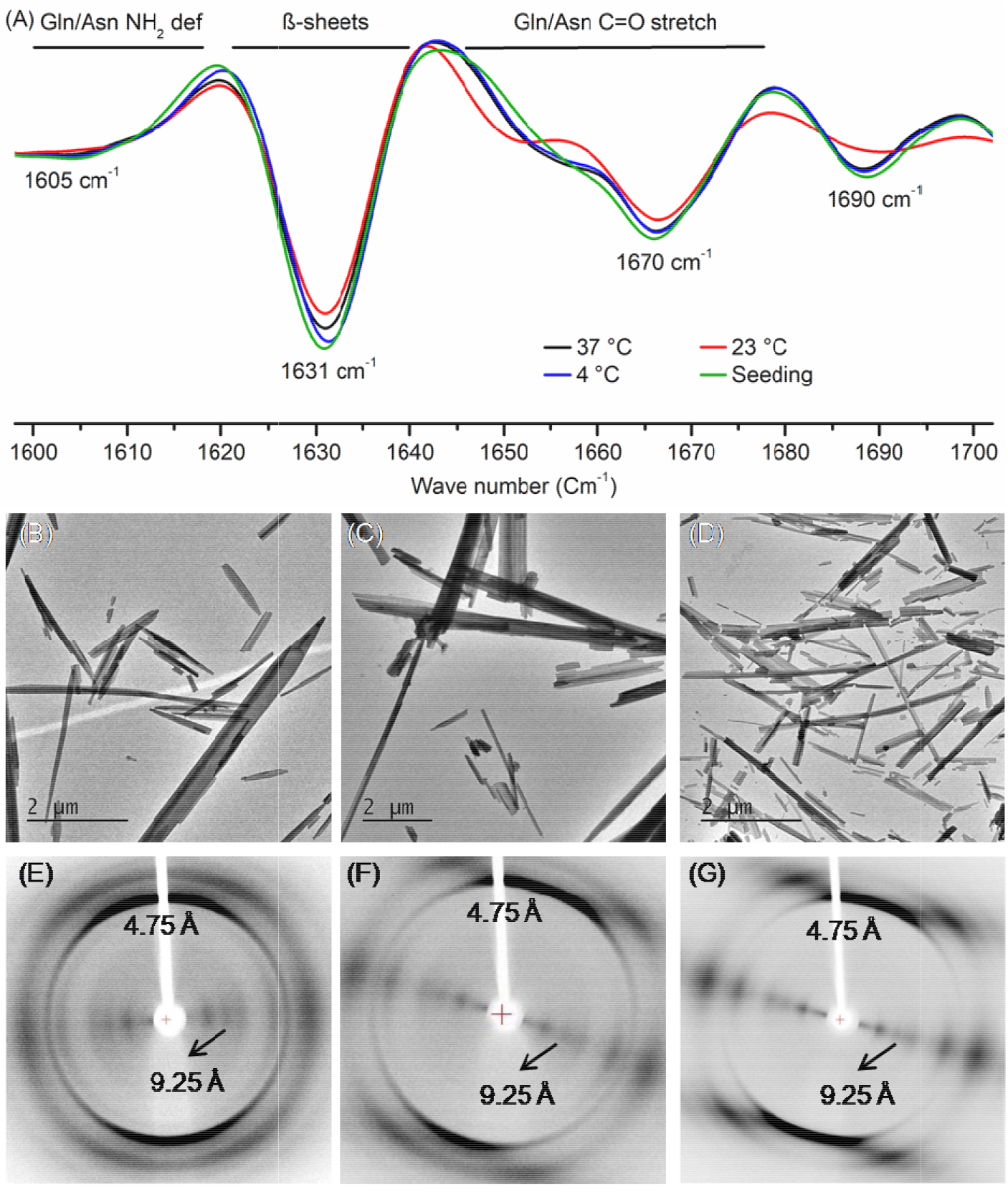
Morphological and structural characterization of mature amyloid fibrils generated at different conditions. (A) Bio-ATR FTIR analysis of normal and seeded aggregates showing characteristic β-sheet bands at 1631 cm^−1^ and 1691 cm^−1^. (B-D)Electron microscopy data showing long unbranched fiber-to-crystalline structures generated at 37 °C (B), 23 °C(C) and 4 °C (D).(E-G) X-ray fiber diffraction images showing the characteristic meridional (4.75 Å) and equatorial (9.25 Å) reflections confirming the cross-β containing amyloid cores for fibers generated at 37 °C (E), 23 °C (F) and 4 °C (G).

**Fig. 5:**
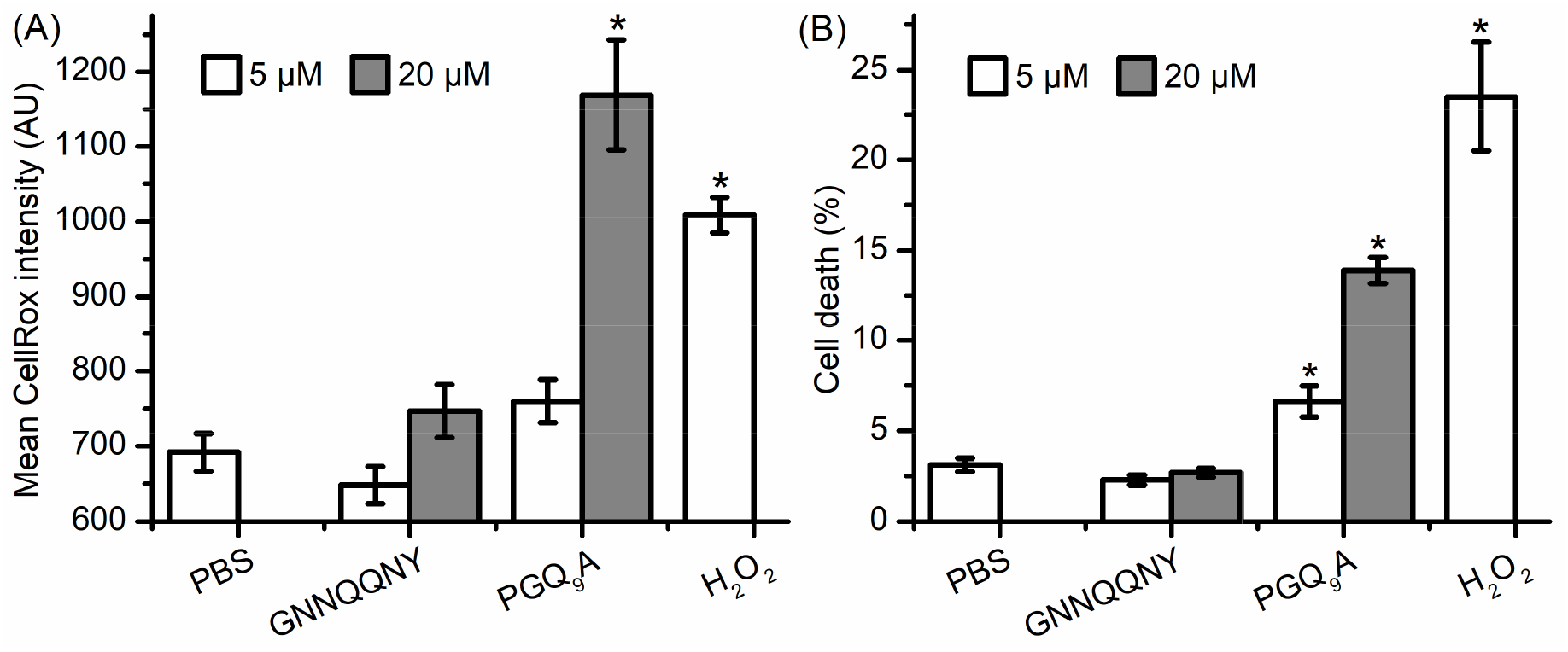
Toxicity data of GNNQQNY in comparison with controls. (A) Oxidative stress exhibited by GNNQQNY and controls upon co-incubating with differentiated SHSY5Y cells for 48 h. (B) Percent cell death observed under similar conditions. Error bars represent the standard deviation calculated from three independent experiments. The statistical significance (*P<0.05) was determined by subjecting the data through one-way ANOVA test.

### Cytotoxicity and cell death analysis

The prion properties of the *Sup*35 protein depends on the glutamine and/or asparagine (QN) rich GNNQQNY sequence that triggers the formation of fibril-core in yeast *Sup*35 prion fibrils.^*3, 10*^ Therefore, we decided to investigate the toxicity of the GNNQQNY fibers using CellRox Green oxidative stress assay and ReadyProbes Live/Dead cell assay. Differentiated SHSY5Y cells were incubated with mature GNNQQNY fibers generated at physiological condition (pH 7.4 and 37 °C). After 48 h of incubation, no difference was observed in the oxidative stress levels and percentage of dead cells in the cells treated with either 5 or 20 μM GNNQQNY aggregates in comparison to PBS-treated cells (Fig. 5A, B). This outcome was compared with that of the fibers formed by a polyglutamine-containing sequence PGQ_9_A and H_2_O_2_ which are known inducers of oxidative stress and cell death.^*52, 53*^ Briefly, the treatment of cells with 5 μM PGQ_9_A for 48 h didn’t induced a significant increase in oxidative stress, however, it resulted in a significant cell death as compared to GNNQQNY and PBS (Fig. 5A, B). Unlike 20 μM GNNQQNY, the 20 μM PGQ_9_A aggregates showed significant increase in oxidative stress and cell death (Fig. 5A, B). As expected, the oxidative stress and cell death induced by H_2_O_2_ was statistically significant as compared to GNNQQNY and PBS (Fig. 5A, B). Overall, these pieces of evidences suggest that the fibers formed by GNNQQNY are non-toxic and induce no cell death.

## Conclusion

Here, we aimed to elucidate the *in vitro* aggregation kinetics of GNNQQNY peptide that was shown to form cross-β containing amyloid structures. This was achieved by using a solubilisation protocol that results in homogenous monomer solution for studying the kinetics. The results are consistent with the nucleation-dependent aggregation kinetics model shown in Fig. 6.^*14*^ In this model, the usually unordered monomers at above their critical concentration interact with each other resulting in the nucleation of β-strands containing species called critical nucleus.^*25*^ The thermodynamic free energy barrier of its formation is generally higher and hence the nucleation event is a highly unfavourable reaction.^*14*^ Hence, the size of the nucleus formed is dependent on both the initial concentration and the temperature of the reaction. Interestingly, the size of the nucleus formed and the length of lag-phase was found to be inversely related to the reaction temperature. The nucleation event is followed by a spontaneous reduction in the monomer concentration and this phase is considered to be the elongation-phase. During this phase, the nucleus acts as a template for further monomer addition leading to the formation of mature amyloid fibers.

**Fig. 6:**
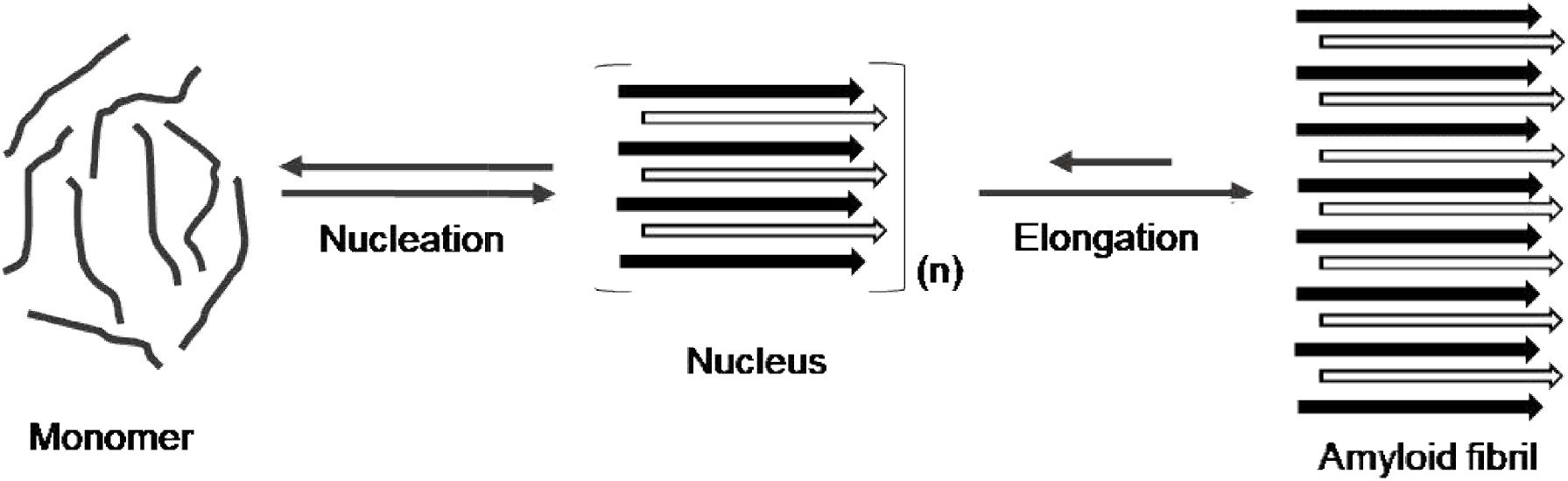
Representative nucleation-dependent aggregation mechanism of GNNQQNY. The unordered GNNQQNY monomers interact with each other through nucleation-phase and results in the formation of a thermodynamically unstable species called nucleus. This nucleus grows spontaneously by further monomer addition leading to the formation of mature amyloid fibers.

Structural characterizations of these fibers confirmed the long unbranched amyloid fibrils rich with cross-β architecture and were found to be similar irrespective of the temperature at which they were grown. In agreement with the literature, the inter-strand and inter-sheet distances within these amyloid fibrils were 4.75 Å and 9.25 Å, respectively.^*14, 34, 36*^ These structures were also stained positive for ThT and CR. Morphological and structural identity of the generated aggregates irrespective of the temperature with that of the published literature validates the conclusions derived in this study. This study further supports the previous theoretical and simulation-based studies through *in vitro* experimental data. The data generated here will provide the basis for further understanding of the kinetics and thermodynamics of GNNQQNY amyloid formation as well as other amyloidogenic protein sequences associated with several systemic and neurodegenerative disorders. The inability of the GNNQQNY aggregates to exhibit oxidative stress and cytotoxicity and the associated stability arising from the β-sheet rich architecture makes them suitable for biotechnological and nanotechnological applications.

## Author contributions

GB conceptualised the study, executed the experiments and interpreted the results. BG and AKT conceptualised the nucleation kinetics analysis, and BG executed and analysed the data. MBM executed the cell work and interpreted the data. Samples for XRD analysis were prepared by GB and MBM, and the data was recorded and analysed by GB and LCS. LCS and AKT supervised the project and managed the finances. Manuscript was prepared by GB and edited by MBM, LCS and AKT.

## Acknowledgement

GB greatly acknowledges the European Molecular Biology Organisation (EMBO) for providing the Short-Term Fellowship award (EMBO-STF 7674) to visit Prof. Louise C. Serpell’s lab at University of Sussex, in the United Kingdom. LCS acknowledges funding from Alzheimer’s Society in support of MBM (AS-PG-16b-010). LCS and MBM are members of the Alzheimer’s Research UK South Coast Network and are grateful for their support. AKT acknowledges Department of Biotechnology (DBT/BSBE/20120020 and DBT/20100368) and Indian Council of Medical Research (ICMR/BSBE/2016489), Govt. of India for the financial support. AKT also acknowledges Shreya Ghosh for her support during nucleation kinetics analysis at 37 °C.

## References

1. Makin, O. S., Atkins, E., Sikorski, P., Johansson, J., and Serpell, L. C. (2005) Molecular basis for amyloid fibril formation and stability, Proc. Natl. Acad. Sci. USA 102, 315–320.

2. Wetzel, R. (2012) Physical chemistry of polyglutamine: intriguing tales of a monotonous sequence, J. Mol. Biol. 421, 466–490.

3. Balbirnie, M., Grothe, R., and Eisenberg, D. S. (2001) An amyloid-forming peptide from the yeast prion Sup35 reveals a dehydrated beta-sheet structure for amyloid, Proc. Natl. Acad. Sci. USA 98, 2375–2380.

4. Reches, M., Porat, Y., and Gazit, E. (2002) Amyloid Fibril Formation by Pentapeptide and Tetrapeptide Fragments of Human Calcitonin, J. Biol. Chem. 277, 35475–35480.

5. Morris, K. L., Rodger, A., Hicks, M. R., Debulpaep, M., Schymkowitz, J., Rousseau, F., and Serpell, L. C. (2013) Exploring the sequence-structure relationship for amyloid peptides, Biochem. J. 450, 275–283.

6. Wickner, R. B., Edskes, H. K., Bateman, D. A., Kelly, A. C., Gorkovskiy, A., Dayani, Y., and Zhou, A. (2013) Amyloids and yeast prion biology, Biochemistry 52, 1514–1527.

7. True, H. L., and Lindquist, S. L. (2000) A yeast prion provides a mechanism for genetic variation and phenotypic diversity, Nature 407, 477–483.

8. Wickner, R. B. (1994) [URE3] as an altered URE2 protein: evidence for a prion analog in Saccharomyces cerevisiae, Science 264, 566–569.

9. Derkatch, I. L., Bradley, M. E., Zhou, P., Chernoff, Y. O., and Liebman, S. W. (1997) Genetic and environmental factors affecting the de novo appearance of the [PSI+] prion in Saccharomyces cerevisiae, Genetics 147, 507–519.

10. Kushnirov, V. V., Vishnevskaya, A. B., Alexandrov, I. M., and Ter-Avanesyan, M. D. (2007) Prion and Nonprion Amyloids, Prion 1, 179–184.

11. Geddes, A. J., Parker, K. D., Atkins, E. D., and Beighton, E. (1968) “Cross-beta” conformation in proteins, J. Mol. Biol. 32, 343–358.

12. Serpell, L. C., Sunde, M., Fraser, P. E., Luther, P. K., Morris, E. P., Sangren, O., Lundgren, E., and Blake, C. C. (1995) Examination of the structure of the transthyretin amyloid fibril by image reconstruction from electron micrographs, J. Mol. Biol. 254, 113–118.

13. Sunde, M., Serpell, L. C., Bartlam, M., Fraser, P. E., Pepys, M. B., and Blake, C. C. (1997) Common core structure of amyloid fibrils by synchrotron X-ray diffraction, J. Mol. Biol. 273, 729–739.

14. Nelson, R., Sawaya, M. R., Balbirnie, M., Madsen, A. O., Riekel, C., Grothe, R., and Eisenberg, D. (2005) Structure of the cross-beta spine of amyloid-like fibrils, Nature 435, 773–778.

15. Sawaya, M. R., Sambashivan, S., Nelson, R., Ivanova, M. I., Sievers, S. A., Apostol, M. I., Thompson, M. J., Balbirnie, M., Wiltzius, J. J., McFarlane, H. T., Madsen, A. O., Riekel, C., and Eisenberg, D. (2007) Atomic structures of amyloid cross-beta spines reveal varied steric zippers, Nature 447, 453–457.

16. Diaz-Avalos, R., Long, C., Fontano, E., Balbirnie, M., Grothe, R., Eisenberg, D., and Caspar, D. L. D. (2003) Cross-beta order and diversity in nanocrystals of an amyloid-forming peptide, J. Mol. Biol. 330, 1165–1175.

17. Nasica-Labouze, J., Meli, M., Derreumaux, P., Colombo, G., and Mousseau, N. (2011) A Multiscale Approach to Characterize the Early Aggregation Steps of the Amyloid-Forming Peptide GNNQQNY from the Yeast Prion Sup-35, PLoS Comput. Biol. 7, e1002051.

18. Nasica-Labouze, J., and Mousseau, N. (2012) Kinetics of Amyloid Aggregation: A Study of the GNNQQNY Prion Sequence, PLoS Comput. Biol. 8, e1002782.

19. Wetzel, R. (2006) Kinetics and thermodynamics of amyloid fibril assembly, Acc. Chem. Res. 39, 671–679.

20. Langkilde, A. E., Morris, K. L., Serpell, L. C., Svergun, D. I., and Vestergaard, B. (2015) The architecture of amyloid-like peptide fibrils revealed by X-ray scattering, diffraction and electron microscopy, Acta Cryst. D 71, 882–895.

21. Thakur, A. K., and Wetzel, R. (2002) Mutational analysis of the structural organization of polyglutamine aggregates, Proc. Natl. Acad. Sci. USA 99, 17014–17019.

22. Burra, G., and Thakur, A. K. (2016) Unaided trifluoroacetic acid pretreatment solubilizes polyglutamine peptides and retains their biophysical properties of aggregation, Anal. Biochem. 494, 23–30.

23. Marshall, K. E., Vadukul, D. M., Dahal, L., Theisen, A., Fowler, M. W., Al-Hilaly, Y., Ford, L., Kemenes, G., Day, I. J., Staras, K., and Serpell, L. C. (2016) A critical role for the self-assembly of Amyloid-β1-42 in neurodegeneration, Sci. Rep. 6, 30182.

24. Burra, G., and Thakur, A. K. (2015) Anhydrous trifluoroacetic acid pretreatment converts insoluble polyglutamine peptides to soluble monomers, Data Brief 5, 1066–1071.

25. Burra, G., and Thakur, A. K. (2019) Insights into the molecular mechanism behind solubilization of amyloidogenic polyglutamine-containing peptides, Pept. Sci. 111, e24094.

26. Maina, M. B., Mengham, K., Burra, G. K., Al-Hilaly, Y. A., and Serpell, L. C. (2020) Dityrosine cross-link trapping of amyloid-β intermediates reveals that self-assembly is required for Aβ-induced cytotoxicity, bioRxiv 007690 [Preprint], doi: https://doi.org/10.1101/2020.1103.1125.007690.

27. O’Nuallain, B., Thakur, A. K., Williams, A. D., Bhattacharyya, A. M., Chen, S., Thiagarajan, G., and Wetzel, R. (2006) Kinetics and Thermodynamics of Amyloid Assembly Using a High-Performance Liquid Chromatography–Based Sedimentation Assay, In Methods Enzymol., pp 34–74, Academic Press.

28. Serpell, L. C., Berriman, J., Jakes, R., Goedert, M., and Crowther, R. A. (2000) Fiber diffraction of synthetic α-synuclein filaments shows amyloid-like cross-β conformation, Proc. Natl. Acad. Sci. USA 97, 4897–4902.

29. Ruiz-Zamora, R. A., Guillaumé, S., Al-Hilaly, Y. K., Al-Garawi, Z., Rodríguez-Alvarez, F. J., Zavala-Padilla, G., Pérez-Carreón, J. I., Rodríguez-Ambriz, S. L., Herrera, G. A., Becerril-Luján, B., Ochoa-Leyva, A., Melendez-Zajgla, J., Serpell, L., and del Pozo-Yauner, L. (2019) The CDR1 and Other Regions of Immunoglobulin Light Chains are Hot Spots for Amyloid Aggregation, Sci. Rep. 9, 3123.

30. Burra, G., and Thakur, A. K. (2017) Inhibition of polyglutamine aggregation by SIMILAR huntingtin N-terminal sequences: Prospective molecules for preclinical evaluation in Huntington’s disease, Pept. Sci. 108, e23021.

31. Saha, I., Singh, V., Burra, G., and Thakur, A. K. (2018) Osmolytes modulate polyglutamine aggregation in a sequence dependent manner, J. Pept. Sci. 24, e3115.

32. Ferrone, F. A. (2006) Nucleation: the connections between equilibrium and kinetic behavior, Methods Enzymol. 412, 285–299.

33. Picken, M. M., and Herrera, G. A. (2015) Thioflavin T Stain: An Easier and More Sensitive Method for Amyloid Detection, In Amyloid and Related Disorders: Surgical Pathology and Clinical Correlations (Picken, M. M., Herrera, G. A., and Dogan, A., Eds.), pp 225–227, Springer International Publishing, Cham.

34. Ostapchenko, V. G., Sawaya, M. R., Makarava, N., Savtchenko, R., Nilsson, K. P., Eisenberg, D., and Baskakov, I. V. (2010) Two amyloid States of the prion protein display significantly different folding patterns, J. Mol. Biol. 400, 908–921.

35. Al-Hilaly, Y. K., Foster, B. E., Biasetti, L., Lutter, L., Pollack, S. J., Rickard, J. E., Storey, J. M. D., Harrington, C. R., Xue, W.-F., Wischik, C. M., and Serpell, L. C. Tau (297-391) forms filaments that structurally mimic the core of paired helical filaments in Alzheimer’s disease brain, FEBS Lett. 594, 944–950.

36. Marshall, K. E., Hicks, M. R., Williams, T. L., Hoffmann, S. V., Rodger, A., Dafforn, T. R., and Serpell, L. C. (2010) Characterizing the Assembly of the Sup35 Yeast Prion Fragment, GNNQQNY: Structural Changes Accompany a Fiber-to-Crystal Switch, Biophys. J. 98, 330–338.

37. Maina, M. B., Bailey, L. J., Wagih, S., Biasetti, L., Pollack, S. J., Quinn, J. P., Thorpe, J. R., Doherty, A. J., and Serpell, L. C. (2018) The involvement of tau in nucleolar transcription and the stress response, Acta Neuropathol Commun 6, 70.

38. Maina, M. B., Bailey, L. J., Doherty, A. J., and Serpell, L. C. (2018) The Involvement of Aβ42 and Tau in Nucleolar and Protein Synthesis Machinery Dysfunction, Front Cell Neurosci 12, 220.

39. Shaw, K. L., Grimsley, G. R., Yakovlev, G. I., Makarov, A. A., and Pace, C. N. (2001) The effect of net charge on the solubility, activity, and stability of ribonuclease Sa, Protein sci. 10, 1206–1215.

40. Kuipers, B. J., and Gruppen, H. (2007) Prediction of molar extinction coefficients of proteins and peptides using UV absorption of the constituent amino acids at 214 nm to enable quantitative reverse phase high-performance liquid chromatography-mass spectrometry analysis, J. Agric. Food. Chem. 55, 5445–5451.

41. Bhattacharyya, A. M., Thakur, A. K., and Wetzel, R. (2005) polyglutamine aggregation nucleation: thermodynamics of a highly unfavorable protein folding reaction, Proc. Natl. Acad. Sci. USA 102, 15400–15405.

42. Kar, K., Jayaraman, M., Sahoo, B., Kodali, R., and Wetzel, R. (2011) Critical nucleus size for disease-related polyglutamine aggregation is repeat-length dependent, Nat. Struct. Mol. Biol. 18, 328–336.

43. Ferrone, F. (1999) Analysis of protein aggregation kinetics, Methods Enzymol. 309, 256–274.

44. Sparrer, H. E., Santoso, A., Szoka, F. C., and Weissman, J. S. (2000) Evidence for the Prion Hypothesis: Induction of the Yeast [*PSI^+^*] Factor by in Vitro-Converted Sup35 Protein, Science 289, 595–599.

45. Groenning, M. (2010) Binding mode of Thioflavin T and other molecular probes in the context of amyloid fibrils-current status, J. Chem. biol. 3, 1–18.

46. Khurana, R., Coleman, C., Ionescu-Zanetti, C., Carter, S. A., Krishna, V., Grover, R. K., Roy, R., and Singh, S. (2005) Mechanism of thioflavin T binding to amyloid fibrils, J. Struct. Biol. 151, 229–238.

47. Rambaran, R. N., and Serpell, L. C. (2008) Amyloid fibrils: abnormal protein assembly, Prion 2, 112–117.

48. Yakupova, E. I., Bobyleva, L. G., Vikhlyantsev, I. M., and Bobylev, A. G. (2019) Congo Red and amyloids: history and relationship, Bioscience Rep. 39, BSR20181415.

49. Glenner, G. G. (1981) The bases of the staining of amyloid fibers: their physico-chemical nature and the mechanism of their dye-substrate interaction, Prog. Histochem. Cytochem. 13, 1–37.

50. Erskine, E., MacPhee, C. E., and Stanley-Wall, N. R. (2018) Functional Amyloid and Other Protein Fibers in the Biofilm Matrix, J. Mol. Biol. 430, 3642–3656.

51. van der Wel, P. C. A., Lewandowski, J. R., and Griffin, R. G. (2007) Solid-State NMR Study of Amyloid Nanocrystals and Fibrils Formed by the Peptide GNNQQNY from Yeast Prion Protein Sup35p, J. Am. Chem. Soc. 129, 5117–5130.

52. Yang, W., Dunlap, J. R., Andrews, R. B., and Wetzel, R. (2002) Aggregated polyglutamine peptides delivered to nuclei are toxic to mammalian cells, Hum. Mol. Genet. 11, 2905–2917.

53. Thakur, A. K., Yang, W., and Wetzel, R. (2004) Inhibition of polyglutamine aggregate cytotoxicity by a structure-based elongation inhibitor, FASEB J. 18, 923–925.

