## Supplementary information for "Nucleation-dependent aggregation kinetics of Yeast *Sup*35 fragment GNNQQNY"

### Standard-curve generation based on analytical Reversed-Phase HPLC

Approximately 1 mg of GNNQQNY peptide was solubilized as previously mentioned in 2.5 mL of milliQ water (pH 2.0), subjected to ultracentrifugation at 80,000 rpm and 25 °C for 2 h and the supernatant was used as stock for determining the standard curve. OD of a series of dilutions prepared from the stock was measured at 220 nm by using a NanoDrop method and the corresponding concentration (µg) was back calculated as per the standard procedure described by Kuipers and Gruppen (2007).[1] Each of these dilutions was passed through Agilent eclipse plus $C_{18}$ column (4.6 mm × 100 mm) connected to Agilent 1260 Infinity Quaternary liquid chromatography system. Gradient of water and acetonitrile containing 0.05% (v/v) TFA at 1 mL/min flow rate was used for elution. The area under the curve monitored at 214 nm for each dilution was plotted against the respective concentration (µg) calculated based on OD to derive the standard-curve that was used for determining the unknown concentration (µg) of the peptide.

The lack of a reproducible solubilisation protocol and the aromatic amino acid residue tryptophan in its sequence makes it practically difficult to study the kinetics of aggregation based on quantitative methods such as analytical reversed-phase high performance liquid chromatography (RP-HPLC). On the other hand, the data generated by qualitative techniques such as Thioflavin T (ThT) and light scattering assays provides no significant practical meaning as they cannot quantify the concentration of the ongoing reaction mixtures more accurately. Here we utilised previous work of Kuirper and Gruppen to develop a method to follow kinetics of aggregation by monitoring the monomer in solution based on a standard curve.[1] The standard curve was generated by plotting the concentration (µg) measured using NanoDrop against the area under the curve observed in chromatogram for different dilutions at 214 nm (Fig. 2A).

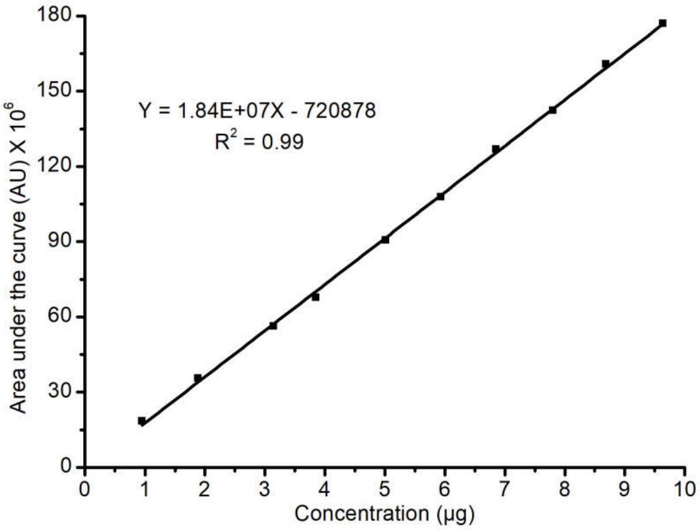


Fig. S1: Standard curve used to determining the unknown concentration

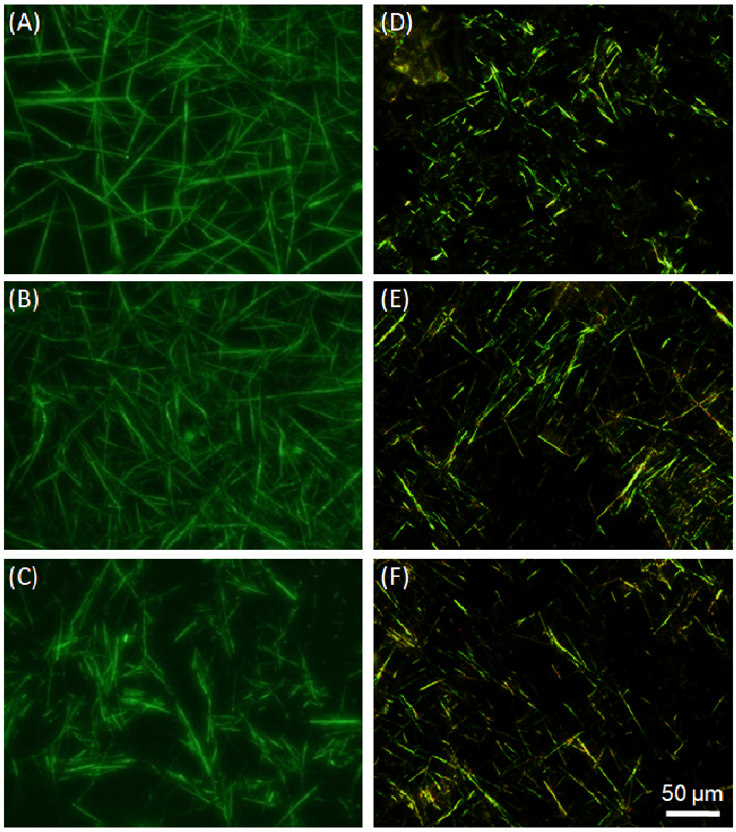

**Fig. S2:** Detection of amyloids based on ThT and CR staining methods. (A-C) Showing the distinct bright *yellow-green* fluorescence of ThT highlighting the amyloid fibrils. (D-F) Congo red stained samples when viewed under cross-polarized light showed characteristic *apple-green* birefringence. (A,D), (B,E) and (C,F) represents the amyloid fibrils generated at 37 °C, 23 °C and 4 °C, respectively.

(1) Kuipers, B. J.; Gruppen, H. *Journal of agricultural and food chemistry* **2007**, *55*, 5445.
